# COMUNET: a tool to explore and visualize intercellular communication

**DOI:** 10.1101/864686

**Authors:** Maria Solovey, Antonio Scialdone

## Abstract

Intercellular communication plays an essential role in multicellular organisms and several algorithms to analyse it from single-cell transcriptional data have been recently published, but the results are often hard to visualize and interpret. We developed COMUNET (Cell cOMmunication exploration with MUltiplex NETworks), a tool that streamlines the interpretation of the results from cell-cell communication analyses. COMUNET uses multiplex networks to represent and cluster all potential communication pathways between cell types. The algorithm also enables the search for specific patterns of communication and can perform comparative analysis between two biological conditions.

To exemplify its use, here we apply COMUNET to investigate cell communication patterns in single-cell transcriptomic datasets from mouse embryos and from an acute myeloid leukemia patient at diagnosis and after treatment. Our algorithm is available on GitHub, along with all the code to perform the analysis reported.

## 1. Introduction

Single-cell RNA sequencing (scRNA-seq) has been extensively used in the last few years to analyse intercellular communication in tissues (see, e.g., (Camp *et al.*, 2017; Puram *et al.*, 2017; Zepp *et al.*, 2017; Zhou *et al.*, 2017; Cohen *et al.*, 2018; Halpern *et al.*, 2018; Skelly *et al.*, 2018; Vento-Tormo *et al.*, 2018; Kumar *et al.*, 2018; Schiebinger *et al.*, 2019; Bonnardel *et al.*, 2019; Sheikh *et al.*, 2019)). Several algorithms to perform these analyses have been published (for instance, (Efremova *et al.*, 2019; Vento-Tormo *et al.*, 2018; Rieckmann *et al.*, 2017; Boisset *et al.*, 2018; Ramilowski *et al.*, 2015) and they all start from a database of interacting partners (e.g., ligand and receptor pairs) to infer, from their expression patterns, a list of potential communication pathways between cell types. Results are visualized with graphs or heatmaps; however, with a large number of potentially communicating cell types and ligand-receptor pairs, these visualization strategies become busy, poorly interpretable and hinder data-driven hypothesis generation.

Here we present COMUNET (Cell cOMmunication exploration with MUltiplex NETworks), a new tool to visualize and interpret cell-cell communication that is based on multiplex networks. COMUNET allows unsupervised clustering of ligand-receptor pairs, search for specific patterns of communication and comparison between two biological conditions, aiding the interpretability of the results and the identification of promising candidate molecules to follow up on.

## 2. Algorithm

### 2.1 Multiplex network

As a first step, COMUNET uses a multiplex network to represent cellular communication pathways (consisting of, e.g., pairs of ligands and receptors) identified by an algorithm of choice (e.g., CellPhoneDB (Vento-Tormo *et al.*, 2018; Efremova *et al.*, 2019)). The input is a set of weight matrices, one for each communication pathway (Figure 1A). As an example, in all the following analyses we apply COMUNET to the output of the CellPhoneDB algorithm (Vento-Tormo *et al.*, 2018; Efremova *et al.*, 2019).

**Figure 1.**
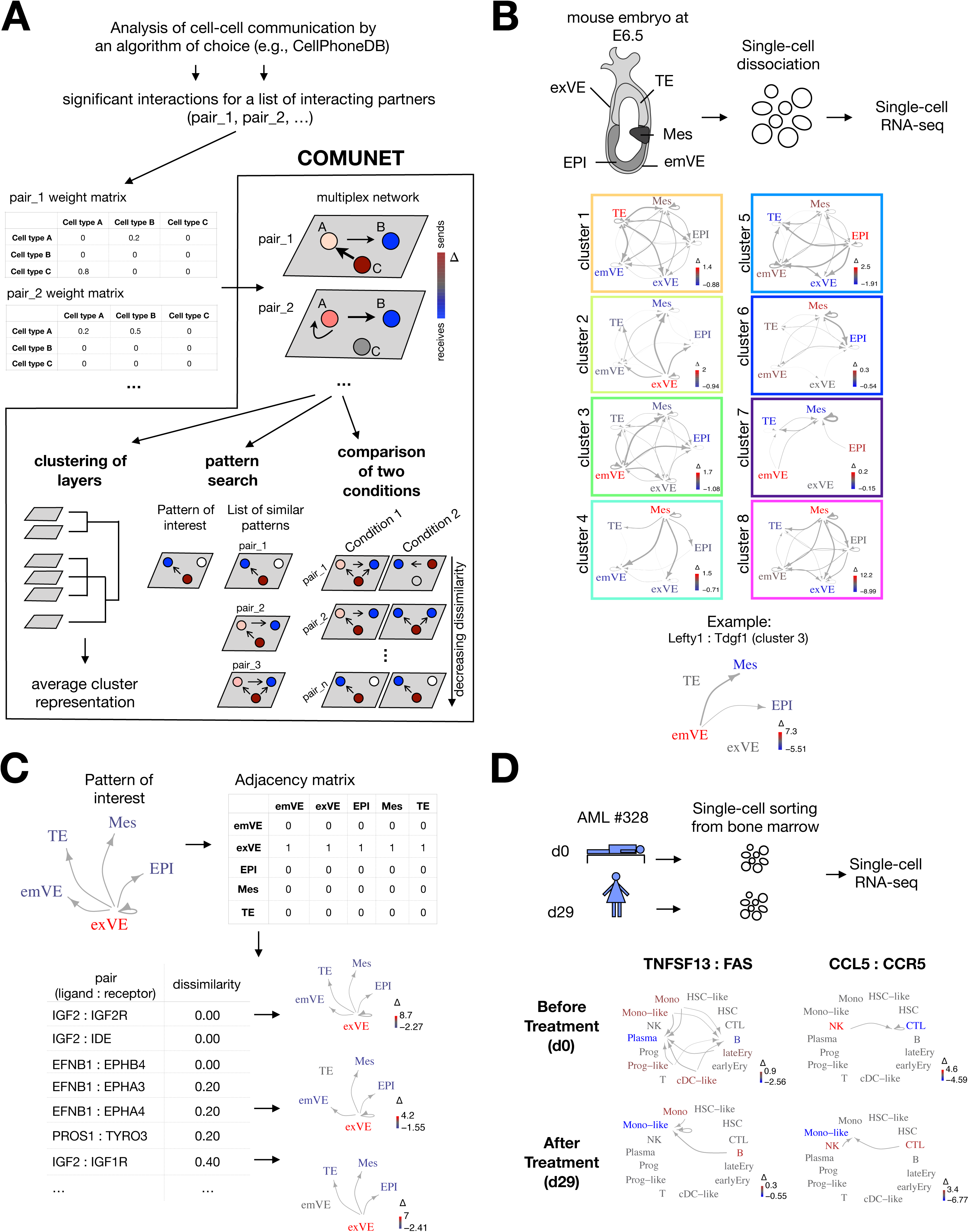
**A.** Schematic representation of COMUNET workflow. Once the weight matrices for a set of interacting partners (e.g., ligands-receptor pairs) are estimated with an algorithm of choice (e.g., CellPhoneDB), COMUNET represents them as layers in a multiplex network, where nodes are cell types (indicated with A, B, C). Each node is colored based on the difference between the weighted in- and out-degree (indicated with Δ), in such a way that the nodes that preferentially send signals are red, while the nodes that preferentially receive signals are blue. Next, COMUNET calculates pairwise dissimilarities between layers in the multiplex network and can perform: clustering of layers, to reveal ligands-receptor pairs sharing similar communication patterns; search of ligands-receptor pairs showing a specific communication pattern; comparison of communication patterns between two biological conditions. **B.** We used COMUNET to identify clusters of ligands-receptor pairs in published scRNA-seq data from E6.5 mouse embryo. The dataset included 5 cell types: extraembryonic visceral endoderm (exVE), trophectoderm (TE), mesoderm (Mes), embryonic visceral endoderm (emVE), and epiblast (EPI). Eight clusters of ligand-receptor pairs were identified, each corresponding to a specific communication pattern, whose average representation is depicted in the squares. In particular, Cluster 3 represents communication from the emVE to EPI and Mes, Lefty1:Tdgf1 being a representative ligand:receptor pair included in this cluster. **C.** As an example of pattern search, using the same scRNA-seq data of panel B, we searched for the ligand-receptor pairs showing the pattern of communication depicted in the top left: i.e., a signal originating from the extraembryonic visceral endoderm and received by all other embryonic tissues, which corresponds to the adjacency matrix shown in the top right. COMUNET returned a list of ligands-receptors sorted by increasing dissimilarity with the specified pattern (bottom left). Bottom right panel illustrates the graphs corresponding to some selected ligand:receptor pairs. **D.** We applied COMUNET to a published scRNA-seq dataset from bone marrow of an AML patient at diagnosis (d0) and after treatment (d29) to find differences in communication patterns between the two time points. TNFSF13:FAS and CCL5:CCR5 are examples of pairs with a dramatic change of communication patterns between d0 and d29 (dissimilarity = 1).

Each weight matrix is interpreted as an adjacency matrix of a directed weighted graph, where the entries are the edge weights. For each node (representing cell types), the difference between the weighted in- and out-degree, Δ, is calculated and encoded in the color of the node. With ligand-receptor pairs, a positive value of Δ indicates that the node (cell type) is mostly communicating by producing and secreting the ligand (“sending” node); conversely, a negative Δ marks nodes that communicate mainly by receiving signals through the receptor (“receiving node”). The arrows of the graph starts at the sending nodes and point to the receiving nodes, while the thickness of an edge indicates the edge weight. The graphs built from the matrices are then stacked together as layers of a multiplex network (Figure 1A).

### 2.2 Dissimilarity measure

Once the ligand-receptor pairs are represented as layers in a multiplex network, COMUNET calculates pairwise dissimilarities between them.

Given two layers α and β, their dissimilarity *d*^α,β^ is defined as:

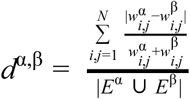

where 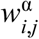 is the weight of the directed edge between nodes *i* and *j* in the layer α, *N* is the number of nodes, *E*^α^ is the set of edges in the layer α, |*E*^α^ ⋃ *E*^β^| is the cardinality of the union of all edges in layers α and β. This definition of dissimilarity can be seen as a modified version of the Jaccard similarity index (Jaccard, 1912) between the sets of edges in the two layers that takes also into account the weights and directionality of the edges.

### 2.3 Clustering

During this step of the analysis, the algorithm identifies clusters of ligand and receptor pairs showing similar patterns of communication (Figure 1A).

A hierarchical clustering of the dissimilarity matrix of layers is performed using the “hclust” R function with the “average” agglomeration method and the results can be visualized as a heatmap or a UMAP plot (McInnes *et al.*, 2018). The number of clusters is estimated using the “cutreeHybrid” R function (package dynamicTreeCut, version 1.63-1(Langfelder *et al.*, 2008)) with “deepSplit” equal to 0 and default “minClusterSize” equal to 6. For each cluster, a graph that represents the “average” pattern is built by averaging the adjacency matrices and the Δ (see section 2.1) of the nodes of all the graphs in the cluster. (Figure 1A, B).

### 2.4 Pattern search

For a more supervised analysis, it is useful to retrieve all ligands and receptor pairs that show a specific communication pattern. To this aim, our algorithm can classify ligand/receptor pairs based on their similarity to a user-specified communication pattern.

A pattern of interest can be specified as a binary adjacency matrix, with sending nodes in the rows and receiving nodes in the columns, 1 indicating the presence of an edge and 0 the absence. The adjacency matrices of the layers are also binarized based on the presence/absence of the edges and the dissimilarities with the user-specified adjacency matrix are calculated (see section 2.2). The output is a list of ligand-receptor pairs sorted by increasing dissimilarity with the user-specified pattern. (Figure 1A and 1C, Suppl Table 2)

### 2.5 Comparative analysis

Given the results of a cell-cell communication analysis of two datasets including the same cell types, COMUNET can estimate the differences in communication patterns between them. First, the results are represented as multiplex networks, as described above; then, pairwise dissimilarities between the shared layers (e.g., layers representing the same pairs of ligands/receptors) of the two datasets are calculated (see Section 2.2). Ligand-receptor pairs are then sorted by decreasing dissimilarity, i.e., from those that change the most across the two datasets to those that remain unaltered (Figure 1A and 1D, Suppl Table 3).

## 3. Applications

Below, we apply COMUNET to scRNA-seq datasets from two different publications to show how the clustering, the pattern search and the comparative analysis can help extract biologically relevant information from a cell-cell communication analysis performed with CellPhoneDB (Vento-Tormo *et al.*, 2018; Efremova *et al.*, 2019).

### 3.1 Mouse embryo

For this example, we used scRNA-seq data from a E6.5 mouse embryo from (Nowotschin *et al.*, 2019) (Figure 1B, Suppl Fig 1). After processing these data with CellPhoneDB, COMUNET finds 8 clusters of ligand-receptor pairs, most of them describing communication from one or two of the cell types (those expressing the ligands) to the other cell types (expressing the receptors). These clusters classify the ligand-receptor pairs according to the pattern of communication they might generate and offer an overview of the main communication patterns present in the data.

As an example, cluster 3 includes signalling originating preferentially from the embryonic visceral endoderm (emVE) to other tissues (Figure 1B, Suppl Fig 1). This cluster includes LEFTY1-TDGF1 (Figure 1B, Suppl Fig 1, Suppl Table 1), which has been described as a visceral endoderm associated signal (Tam and Loebel, 2007).

In addition to this unsupervised analysis, all ligand/receptor pairs having a specific communication pattern can be identified. For example, in Figure 1C and Suppl Table 2, we show how a list of all ligands-receptors that might be responsible for signalling from the extra-embryonic visceral endoderm to all the other cell types can be easily obtained.

### 3.2 Acute Myeloid Leukemia (AML) patient

As a further example, we used the data from (van Galen *et al.*, 2019) to show how COMUNET can find differences in communication patterns between two datasets. This publication includes a scRNA-seq data of a bone marrow sample of a 74 years old newly diagnosed female AML patient (#328) at diagnosis (day 0) and after Azacitidine-Venetoclax-C1D27 treatment (day 29). We compared the communication patterns in the two samples and revealed that 6 ligand-receptor pairs dramatically change their pattern of communication upon treatment. Among those we found TNFSF13-FAS and CCL5-CCR5, for which both the ligand and the receptor play a role in AML and other cancers (Chapellier *et al.*, 2019; Gmeiner *et al.*, 2015; Brenner *et al.*, 2016; Mollica Poeta *et al.*, 2019; Aldinucci and Colombatti, 2014) (Figure 1D, Suppl Fig 2, Suppl Table 3)

### Conclusions

COMUNET is a powerful tool for visualization and analysis of intercellular communication. It can be applied to the output of virtually any algorithm that analyzes cellular communication from any kind of data (not only single-cell transcriptomics), to help explore the results and extract biologically relevant information from them.

The code and a fully commented tutorials to reproduce the analyses and the figures included in this paper are freely available from GitHub at this link: https://github.com/ScialdoneLab/COMUNET.

## Acknowledgments

We thank Jonathan Fiorentino, Gabriele Lubatti and Frank Ziemann for useful discussions and comments on the tutorials.

---

**Supplemental Figure 1.**
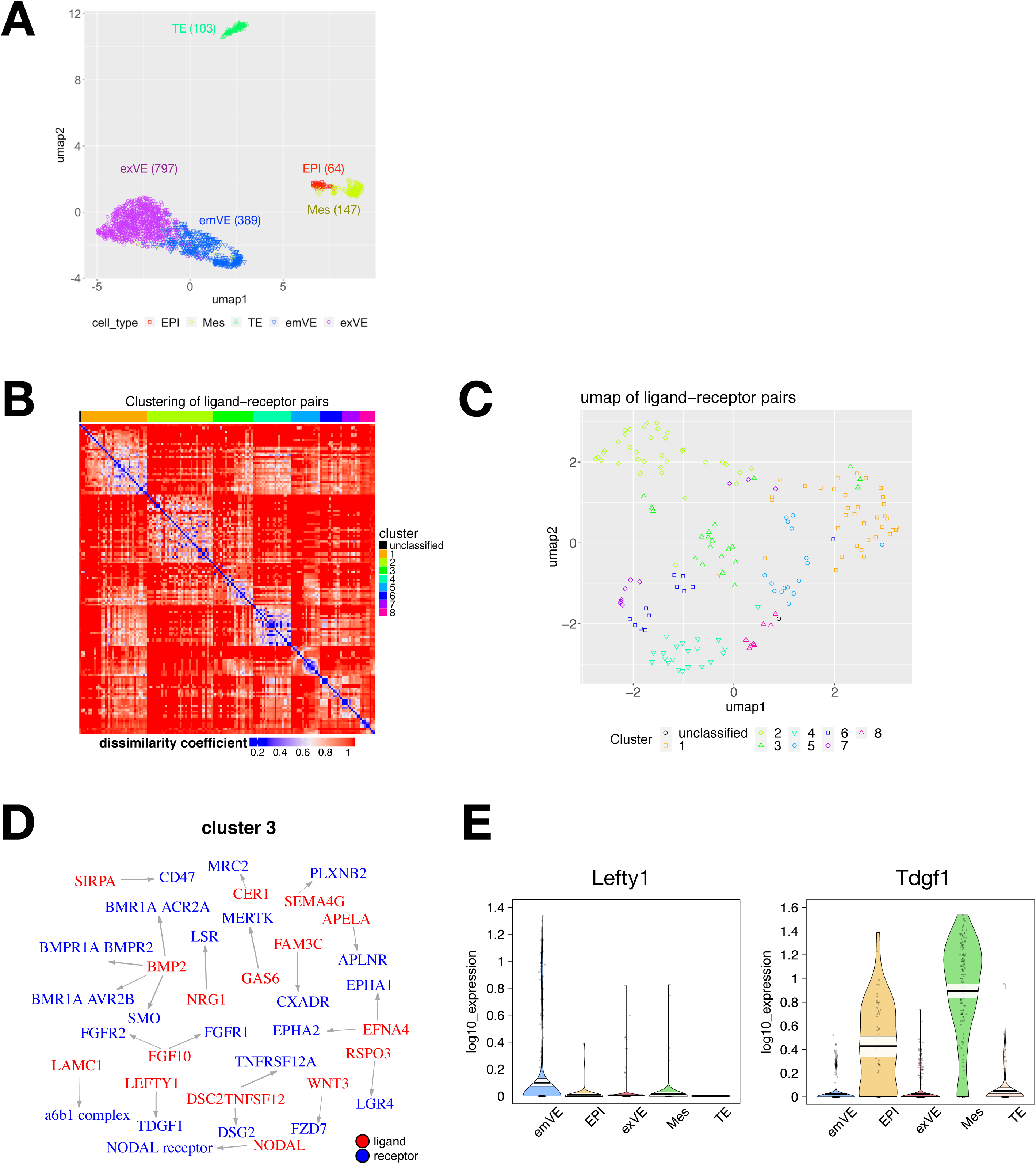
E6.5 mouse embryo data. A. UMAP of the E6.5 mouse embryo scRNA-seq data (supset to 1500 cells). Cell types are: embryonic visceral endoderm (emVE), epiblast (EPI), extraembryonic visceral endoderm (exVE), mesoderm (Mes), and trophectoderm (TE). B. Heatmap showing the dissimilarity of ligand:receptor pairs in the scRNA-seq data from E6.5 mouse embryo (supset to 1500 cells). Rows and columns are sorted by cluster number, the color represents dissimilarity between two pairs with 0 for identical pattern (blue), 1 for completely different pattern (red). C. UMAP of ligand-receptor pairs colored by cluster. D. Ligand and receptors pairs in cluster 3. Ligands are represented in red, receptors in blue; arrows go from the ligand to the associated receptor. E. Log10 expression levels of Lefty1 and Tdgf1.

**Supplemental Figure 2.**
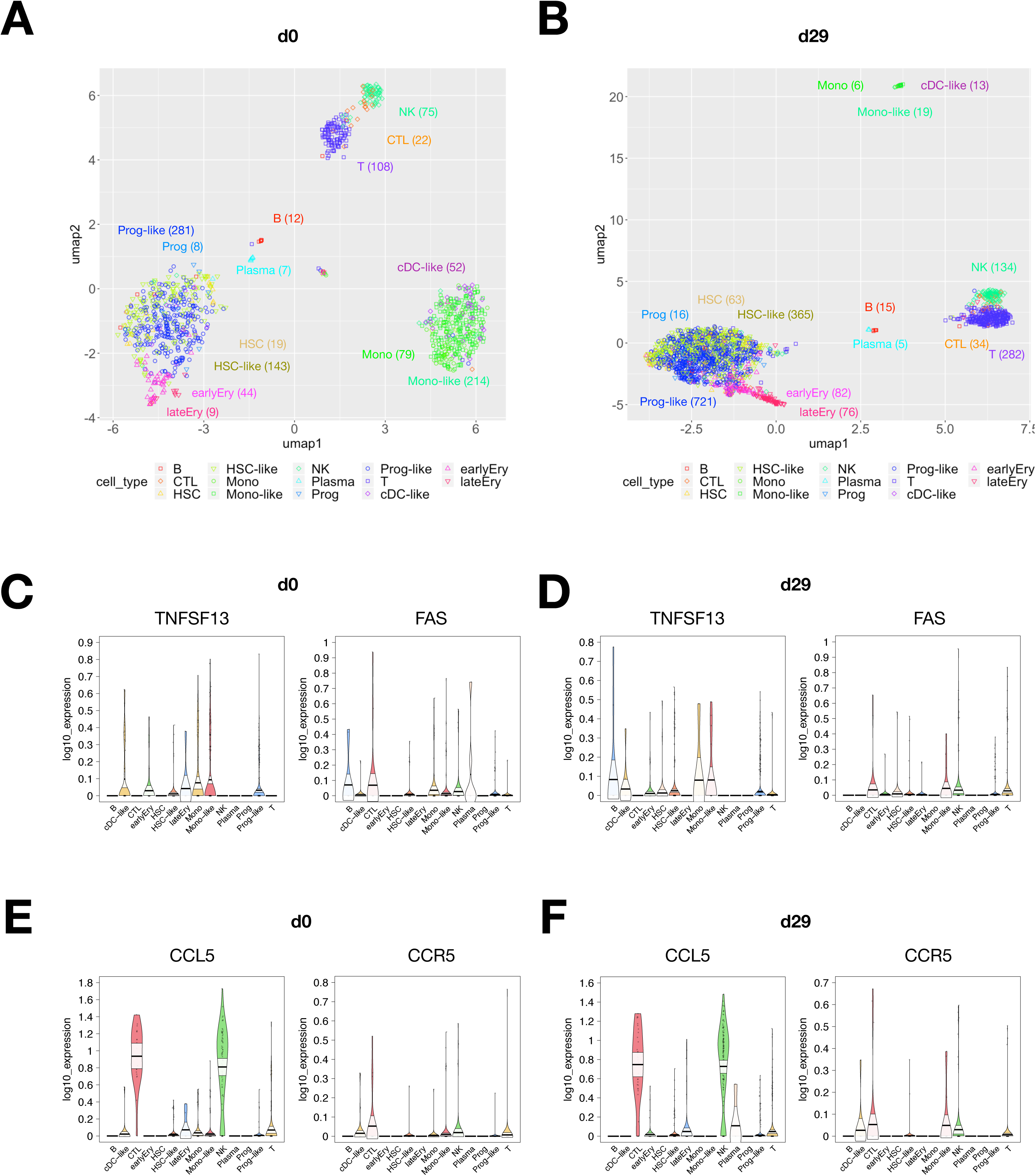
AML patient data. A-B. UMAP of bone marrow of AML patient at d0 (panel A) and d29 (panel B). Number of cells in each population is represented in brackets. C-D. Log10 expression levels of TNFSF13 and FAS at d0 (panel C) and d29 (panel D). E-F. Log10 expression levels of CCL5 and CCR5 at d0 (panel E) and d29 (panel F).

## Notes

https://github.com/ScialdoneLab/COMUNET

## References

Aldinucci,D. and Colombatti,A. (2014) The inflammatory chemokine CCL5 and cancer progression. Mediators Inflamm., 2014, 292376.

Boisset,J.-C. et al. (2018) Mapping the physical network of cellular interactions. Nat. Methods, 15, 547–553.

Bonnardel,J. et al. (2019) Stellate Cells, Hepatocytes, and Endothelial Cells Imprint the Kupffer Cell Identity on Monocytes Colonizing the Liver Macrophage Niche. Immunity, 51, 638–654.e9.

Brenner,A.K. et al. (2016) A Subset of Patients with Acute Myeloid Leukemia Has Leukemia Cells Characterized by Chemokine Responsiveness and Altered Expression of Transcriptional as well as Angiogenic Regulators. Front. Immunol., 7, 205.

Camp,J.G. et al. (2017) Multilineage communication regulates human liver bud development from pluripotency. Nature, 546, 533.

Chapellier,M. et al. (2019) Arrayed molecular barcoding identifies TNFSF13 as a positive regulator of acute myeloid leukemia-initiating cells. Haematologica, 104, 2006–2016.

Cohen,M. et al. (2018) Lung Single-Cell Signaling Interaction Map Reveals Basophil Role in Macrophage Imprinting. Cell, 175, 1031–1044.e18.

Efremova,M. et al. (2019) CellPhoneDB v2.0: Inferring cell-cell communication from combined expression of multi-subunit receptor-ligand complexes. bioRxiv, 347.

van Galen,P. et al. (2019) Single-Cell RNA-Seq Reveals AML Hierarchies Relevant to Disease Progression and Immunity. Cell, 176, 1265–1281.e24.

Gmeiner,W.H. et al. (2015) Thymineless death in F10-treated AML cells occurs via lipid raft depletion and Fas/FasL co-localization in the plasma membrane with activation of the extrinsic apoptotic pathway. Leuk. Res., 39, 229–235.

Halpern,K.B. et al. (2018) Paired-cell sequencing enables spatial gene expression mapping of liver endothelial cells. Nat. Biotechnol., 36, 962–970.

Jaccard,P. (1912) THE DISTRIBUTION OF THE FLORA IN THE ALPINE ZONE.1. New Phytol., 11, 37–50.

Kumar,M.P. et al. (2018) Analysis of Single-Cell RNA-Seq Identifies Cell-Cell Communication Associated with Tumor Characteristics. Cell Rep., 25, 1458–1468.e4.

Langfelder,P. et al. (2008) Defining clusters from a hierarchical cluster tree: the Dynamic Tree Cut package for R. Bioinformatics, 24, 719–720.

McInnes,L. et al. (2018) UMAP: Uniform Manifold Approximation and Projection. JOSS, 3, 861.

Mollica Poeta,V. et al. (2019) Chemokines and Chemokine Receptors: New Targets for Cancer Immunotherapy. Front. Immunol., 10, 379.

Nowotschin,S. et al. (2019) The emergent landscape of the mouse gut endoderm at single-cell resolution. Nature, 569, 361–367.

Puram,S.V. et al. (2017) Single-Cell Transcriptomic Analysis of Primary and Metastatic Tumor Ecosystems in Head and Neck Cancer. Cell, 171, 1611–1624.e24.

Ramilowski,J.A. et al. (2015) A draft network of ligand-receptor-mediated multicellular signalling in human. Nat. Commun., 6, 7866.

Rieckmann,J.C. et al. (2017) Social network architecture of human immune cells unveiled by quantitative proteomics. Nat. Immunol., 18, 583–593.

Schiebinger,G. et al. (2019) Optimal-Transport Analysis of Single-Cell Gene Expression Identifies Developmental Trajectories in Reprogramming. Cell, 176, 928–943.e22.

Sheikh,B.N. et al. (2019) Systematic Identification of Cell-Cell Communication Networks in the Developing Brain. iScience, 21, 273–287.

Skelly,D.A. et al. (2018) Single-Cell Transcriptional Profiling Reveals Cellular Diversity and Intercommunication in the Mouse Heart. Cell Rep., 22, 600–610.

Tam,P.P.L. and Loebel,D.A.F. (2007) Gene function in mouse embryogenesis: get set for gastrulation. Nat. Rev. Genet., 8, 368–381.

Vento-Tormo,R. et al. (2018) Single-cell reconstruction of the early maternal-fetal interface in humans. Nature, 563, 347–353.

Zepp,J.A. et al. (2017) Distinct Mesenchymal Lineages and Niches Promote Epithelial Self-Renewal and Myofibrogenesis in the Lung. Cell, 170, 1134–1148.e10.

Zhou,J.X. et al. (2017) Extracting Intercellular Signaling Network of Cancer Tissues using Ligand-Receptor Expression Patterns from Whole-tumor and Single-cell Transcriptomes. Sci. Rep., 7, 8815.

